# Aquatic plant mycobiomes change with watershed urbanization

**DOI:** 10.1101/2024.06.23.600282

**Authors:** Jacob Mora, Matt Olson, Sara S. Rocks, Geoffrey Zahn

## Abstract

Urban expansion, projected to triple globally from 2000 to 2030, significantly impacts biodiversity and ecosystem processes, including those of microbial communities. While extensive research has examined urbanization effects on macro-organisms, the impacts on microbial communities, particularly those associated with aquatic plants, remain underexplored. This study investigates the fungal endophyte communities in the pollution-tolerant aquatic plant Ranunculus aquatilis along an urbanization gradient in the Provo River, Utah, USA, a rapidly urbanizing region. We collected plant and adjacent water samples from ten locations along the river, spanning from rural to urbanized areas, and conducted DNA amplicon sequencing to characterize fungal community composition.

Our results show a significant decline in fungal alpha diversity downstream, correlated with increased urbanization metrics such as impervious surface area and developed land cover. Specifically, fungal ASV richness and Shannon diversity decreased as urbanization intensified, driven primarily by a reduction in rare taxa. Despite a stable core microbiome dominated by a few taxa, the overall community structure varied significantly along the urban gradient, with notable shifts in dominant fungal taxa. Contrary to expectations, no detectable levels of heavy metals were found in water samples, suggesting that other urbanization-related factors, potentially including organic pollutants or plant stress responses, influence fungal endophyte communities.

Our findings underscore the need for further investigation into the mechanisms driving these patterns, particularly the roles of organic pollution, nutrient loads, and plant stress. Understanding these interactions is crucial for predicting the impacts of continued urbanization on freshwater ecosystems and their associated microbial communities.

## Introduction

Global urban landscapes are projected to triple in area from 2000 to 2030. Although urban expansion is often treated as a local issue, this aggregate predicted global increase in urban area will have profound environmental effects (Seto et al. 2012). A wealth of research demonstrates that the impacts of urbanization on both biodiversity and ecosystem processes can be profound and sudden. For example, urbanization has dramatic effects on plant communities (de Barros Ruas et al. 2022), soil microbial communities (Liu et al. 2022), and their associated ecosystem service values (Liu et al. 2022).

The vast majority of studies on urbanization and biodiversity has focused on macro-organisms, so there is a great need to expand and clarify predictions of how urbanization will affect crucially important microbial communities as well (Li et al. 2010; Liu et al. 2022). Some progress in this area has begun to show that microbial taxa are just as strongly influenced by urbanization pressure as macro-organisms. For instance, urbanization has been correlated with large changes to free-living aquatic fungal community structure, as well as animal microbiomes (Murray et al. 2020; Yuan et al. 2020; Numberger et al. 2022).

The exact mechanisms for the role of urban conversion impacting microbial biodiversity are still unclear and likely vary with location, but potential explanations include increased metal and organic pollution, land-use shifts that result in breaks from historical nutrient cycles, and local climate impacts (Bai et al. 2017). These effects are typically first detected in, and have amplified impacts on, watersheds within a given ecosystem (Lewis et al. 2007; Aitkenhead-Peterson et al. 2011; Blaszczak et al. 2019). Aquatic invertebrates, for example, are particularly sensitive to even small increases in urbanization (Snyder and Young 2020), and metals such as Zn, Cu, Ni, and Cd can have strongly negative effects on growth and sporulation of some aquatic fungi (Azevedo and Cássio 2010). This potential for early detection and outsized impacts makes riverine systems an excellent model for studying the geographic structure of communities in relation to varying urbanization pressure.

Riverine systems have been successfully utilized to show that free-living bacteria and fungi are profoundly affected by increases in urban land-use. These free-living taxa are largely saprotrophic and dependent on organic inputs from riparian vegetation. Bacterial communities in a stream water column can be altered by decreased carbon and increased nitrogen inputs (Wang et al. 2011; Hosen et al. 2017) due to urbanization. Shifts in community structure of free-living fungal communities has also been linked to urbanization, with those shifts being tied to pollution and decreased dissolved oxygen (Medeiros et al. 2009; Moreirinha et al. 2011; Li et al. 2023).

Though it is clear that free-living aquatic microbes are impacted by known urbanization factors, it is uncertain whether host-associated microbes experience the same outcomes. These taxa, such as endophytic fungi, may remain largely protected by their hosts from small shifts in water chemistry. Though aquatic plant microbiomes have been mostly overlooked in favor of terrestrial plants (Sandberg et al. 2014), they are likely providing similar services and may even provide their hosts with resilience to urbanization pressures. Investigating the effects of watershed urbanization on aquatic plant microbiomes will be crucial for predicting the outcomes of projected urbanization on freshwater systems.

To characterize the effect of increasing urbanization on host-associated microbes, we assessed fungal meta-amplicon data from the pollution-tolerant aquatic plant, *Ranunculus aquaticus*, along an increasing urbanization gradient in the Provo River, UT, USA. This watershed is located in Utah County, which is noted to be among the fastest growing urban centers of the United States (US Census Bureau, 2020), and exists along an urbanization gradient with a sharp boundary between low and high urban land-use. We collected plant tissue samples along this gradient and characterized the fungal endophyte community along with water column pollution measures known to affect free-living aquatic fungi in an effort to quantify these effects in host-associated fungi.

## Methods

### Research area and sampling

The Provo River connects the Jordanelle Reservoir to the Utah Lake in northern central Utah. The climate of the region is semi-arid, with an average high temperature in July of 32 C and an average low temperature in January of –7 C. The topography the river traverses is from the high mountainous area of the Wasatch Front Range for 114 km to its terminus at Utah Lake. Our sampling area covered 15 km of the river near its end (Fig. 1b). Ten locations were chosen along the river. Although each sampling location moving downstream from the first location exists along a continuous gradient of urbanization, five locations were upstream of the major metropolitan area and five were within, giving a strong increase in urbanization between the first five and last five sites (Fig. 1a). At each of the sampling locations three samples of *R. aquatilis* were taken haphazardly from different plant patches. Plant samples averaged around 5 cm and included sections of the root shoot and leaf of the plant. Water column samples were also taken adjacent to each sampled plant and stored in 50mL Falcon tubes. All samples were transported to the lab on ice and refrigerated for up to 24 hours before processing.

**Figure 1.**
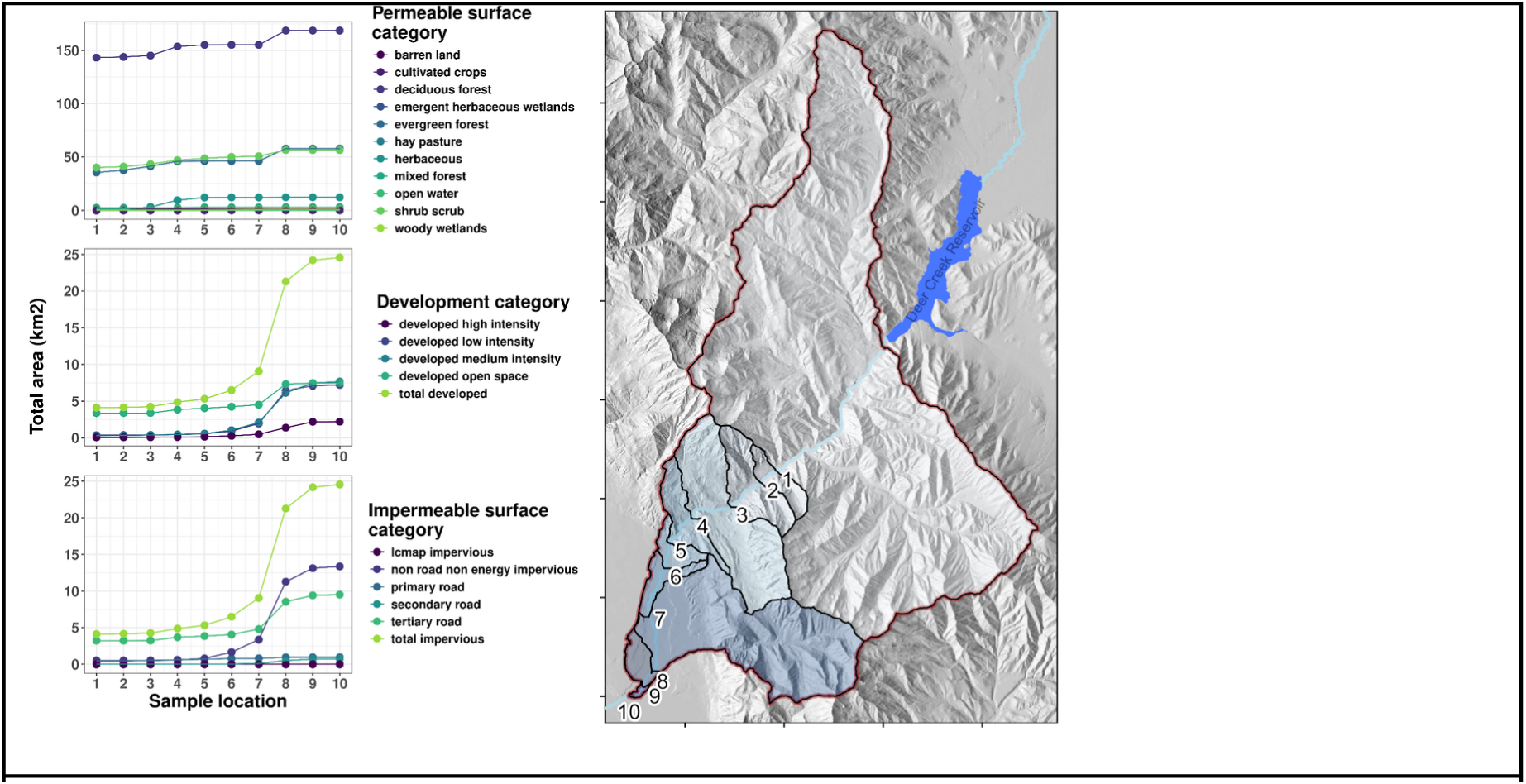
– Watershed map and land use at each sampling location. Left panel: Total area (km2) of land use categories for permeable, developed, and impermeable surface. Colors indicate subcategories of each surface cover. Right panel: Map of watershed at each numbered sampling location.

Plant tissues were surface-sterilized in a two-step wash of 10% bleach (1 min), 70% ethanol (1 min), and then rinsed in 3 changes of sterile water. After surface sterilization, 0.25g of fresh plant roots and shoots were subsampled and frozen for DNA extraction.

### DNA extraction and amplicon sequencing

DNA extraction was performed on all plant samples as well as two negative controls (sterile PCR water) using the DNEasy PowerSoil Pro Kit (Qiagen, MD, USA) according to manufacturer instructions. Genomic DNA from samples and negative controls was amplified with the ITS1F (CTTGGTCATTTAGAGGAAGTAA) (Gardes and Bruns 1993) and ITS2 (GCTG CGTTCTTCATCGATGC) (White et al. 1990) primers modified with the addition of Illumina adaptors (Caporaso et al. 2011) using the following protocol: 98 °C 2 min; 22 cycles of 98 °C 15 s, 52 °C 30 s, 72 °C 30 s; 72 °C 2 min). After 22 cycles, the PCR product was diluted 1:12 and 1 mL of this was used as a template for 8 more rounds of PCR with a 60 °C annealing temperature in which bidirectional barcodes bound to reverse-complemented Illumina adaptors acting as primers. Resulting barcoded libraries were cleaned, normalized, and sequenced on a full Illumina MiSeq run (V3 chemistry, 2 × 300 bp) by Novogene (Sacramento, CA, USA).

### Bioinformatics

The full bioinformatics pipeline, including all tool parameters and statistical methods, can be accessed in the archived GitHub repository associated with this manuscript (Geoffrey Zahn 2023). The ITS1 region was extracted from all amplicon sequences using ITSxpress (Rivers et al. 2018). Reads were then quality screened and reverse reads were discarded (Pauvert et al. 2019). Forward reads were processed in R using the *DADA2* package (Callahan et al. 2016) to remove any reads with uncalled bases and to truncate reads where quality scores dropped significantly. Quality-filtered reads were then used to estimate and correct sequencing errors, and remove de novo-detected chimeras within the *DADA2* package. Contaminant sequences found in negative controls were removed using the prevalence method in the *decontam* R package (Davis et al. 2018). Cleaned and filtered ASVs were then assigned taxonomy with the RDP Classifier algorithm (Wang et al. 2007) against the UNITE_Euk database v. 9.0 (Abarenkov et al. 2023). Any ASVs matching non-fungal taxa, and those that only matched to “Kingdom Fungi” with no more specific taxonomic fungal affiliation, were removed. The remaining ASVs that were taxonomically assigned as fungi were used in all downstream analyses within the *phyloseq* R package (McMurdie and Holmes 2013). Assigned fungal names were checked against the MycoBank synonym database using the *mycobank* R package (Geoffrey Zahn 2024) to assure the validity of names and update any older synonyms in the UNITE database with currently accepted taxonomy. Absolute ASV counts from each sample were transformed to relative abundance values to account for the compositional nature and sequence heterogeneity inherent in Illumina datasets (Gloor et al. 2017).

### Fungal guild analyses

Fungal ASVs were assigned to putative guilds based on assigned taxonomy (Nguyen et al. 2016) using the *FunGuildR* R package (Furneaux and Song 2023). Each ASV was only assigned a major guild of saprotroph, mutualist, or pathogen if confidence was ‘probable’ or higher. Fungal ASVs that were not assigned unambiguous taxonomy at at least the genus level were not assigned a major guild type.

### Urbanization and land use measurement

Impervious surface area was calculated for the individual watershed catchment of each sampling site to better understand the influence of urbanization. Watershed delineation was performed using WhiteboxTools, an open-source R package developed at the University of Guelph’s Geomorphometry and Hydrogeomatics Research Group (Wu and Brown 2022), and a 10-meter resolution digital elevation model (DEM) from the USGS 3D Elevation Program (Dewitz 2023). The delineated watersheds were then used with classified Landsat satellite imagery from the 2021 National Land Cover Database (NLCD) program to calculate the impervious surface area for each unique drainage basin. The NLCD imperviousness products represent urban impervious surfaces (developed) over every 30-meter pixel in the continental United States. The NLCD also includes an impervious surface descriptor layer that identifies types of roads, buildings, and energy production sites to enable detailed analysis of developed features.

### Water chemistry

Soluble metal concentrations (Cr, Co, Ni, Cu, Zn, and Cd) were determined using an Agilent 4210 microwave plasma-atomic emission spectrometer (MP-AES) coupled with an SPS-4 autosampler and nitrogen generator (Agilent Technologies, Santa Clara, CA, USA). The atomic wavelengths for analysis were selected for maximum signal intensity and were 425.433 nm for Cr, 340.512 nm for Co, 352.454 nm for Ni, 323.754 nm for Cu, 213.857 nm for Zn, and 228.802 nm for Cd. Few interferences were expected at these wavelengths. Six standards were prepared from a certified, mixed-metal standard stock solution (5 mg/L in 5% nitric acid, Agilent Technologies) over a concentration range of 0.10 ppm to 5.0 ppm. Correlation coefficients were all greater than 0.999. A calibration blank of 2% nitric acid was prepared using trace-metal grade nitric acid (Fisher Scientific) and 18.2 MΩ cm water. All samples and standards were measured with three replicates, a pump speed of 15 rpm, a 15 s stabilization time, a 3 s read time and a 30 s rinse time.

### Statistical analyses

For alpha diversity measures, generalized linear regression models were fit for all outcomes using the proportion of developed and impervious land in the watershed, along with all development classifications and sampling location as interactive predictors. These full models were simplified using stepwise AIC model selection with the MASS R package (Venables and Ripley 2010), and models were refit using the selected predictors and location. Sample location was included in all models, regardless of AIC selection outcomes, in order to account for the fact that location is inherently tied to urbanization in this system (see Fig. 1). The effect of urbanization metrics on beta-diversity was tested with permutational ANOVA using the *vegan* R package and a Bray-Curtis dissimilarity metric (Okansen et al. 2016). Variable correlation tests were conducted with Pearson’s product-moment correlation in R. Multiple regression on matrices was performed with the *ecodist* R package version 2.1.3 (Goslee and Urban 2007).

The core microbiome was defined as taxa with a relative abundance greater than 1% in at least 50% of samples. Rare taxa were defined as those having less than 0.1% relative abundance total across all samples.

## Results

### Sequencing and community structure

After quality filtration, 429,592 high quality reads remained for all downstream applications, resulting in 299 fungal ASVs. The mean read count per sample was 17,900. The majority of fungal ASVs (79%) were assigned to Ascomycota, followed by Basidiomycota (12%) and 1% or less were assigned to each of Blastocladiomycota, Chytridiomycota, Glomeromycota, Mortierellomycota, Olpidiomycota, and Rozellomycota (S.I. Fig 1).

### Urbanization increased sharply along the watershed

The Provo River flows southwest from rural upland Provo Canyon into the urbanized Utah Valley. Undeveloped portions of the watershed are comprised mostly of deciduous forest, scrublands, and coniferous forest, with the total area of each gradually increasing as the watershed progressively grew downstream. Other undeveloped (permeable) surface types, including herbaceous cover, remain consistently low throughout the watershed. Along the river corridor, watershed impervious surface area increases from 4.12 km2 at the initial sampling site in the canyon (Site 1) to 24.58 km2 at the final site in the valley (Site 10), reflecting a much greater urban influence further downstream.

The rise in impervious surface area is initially gradual along the river corridor, but accelerates sharply below Site 6, marking the transition from a rural to an urbanized watershed. The rapid change in land use from undeveloped to urban coincided with the Provo River’s entry into residential and commercial developments that adjoin the Wasatch Front mountain range. The bulk of the rapid increase in developed and impermeable surface was driven by tertiary roads and non-road, non-energy impermeable surfaces including parking lots, driveways, and rooftops (Figure 1a).

### Heavy metals were too low to detect

None of the water samples from any site had detectable levels of soluble metals (Cr, Co, Ni, Cu, Zn, and Cd). The instrument detection threshold is between 100 and 500 ppb, and each sample was measured in triplicate. Consequently, no quantifiable data on metal concentrations were obtained in this study. Further, our pH measurements of water samples varied from 6.8 to 7.1 along the river, with no consistent pattern.

### The core microbiome was dominated by a few taxa

The core microbiome, designated as taxa that showed at least 1% relative abundance in at least 50% of all samples, was comprised of ten fungal ASVs. The nine most abundant of the core members belonged to Ascomycota, and one member (*Malasseziaceae sp.*) belonged to Basidiomycota. The core ASVs were assigned to a mix of endophyte, plant pathogen, and saprotroph guilds (SI Table 1).

The core membership was dominated by several highly abundant taxa. *Plectosphaerella cucumerina, Lemonniera centrosphaera, Neopyrenochaeta annellidica,* and *Neonectria candida* comprised an average of 48% of reads among all samples throughout the watershed, and greater than 80% of reads in more than half of the sampled locations. The composition of the core taxa shifted near the urban transition zone. While all these taxa were present with at least 1% relative abundance in the majority of all sampling locations, their relative abundance values varied. For example, *N. candida* made up a greater proportion of reads in the upper half of the watershed, while *Cladosporium subuliforme* was a more dominant community member in the lower urbanized portion of the watershed (Figure 2).

**Figure 2.**
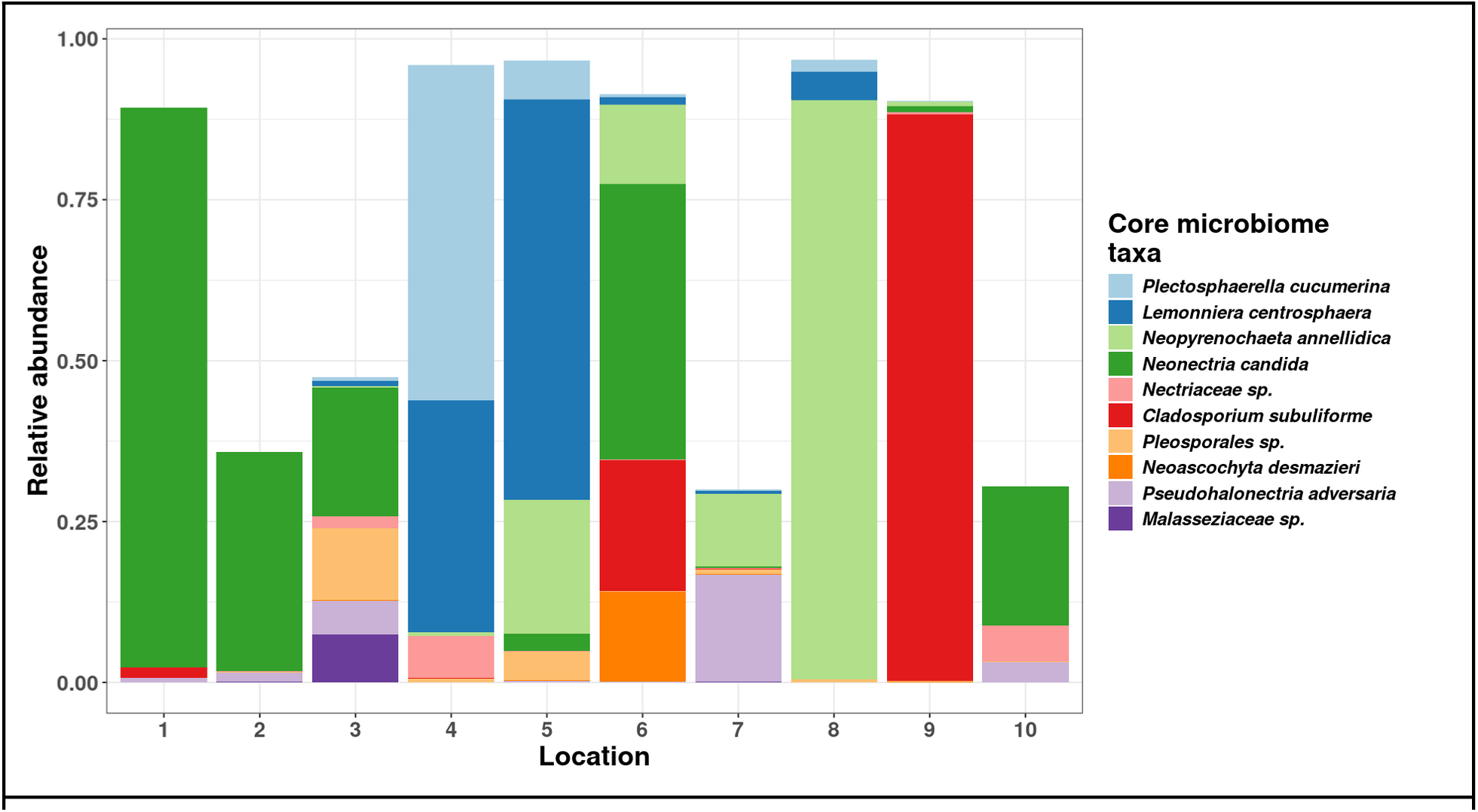
– Total relative abundance of the ten core microbiome taxa at each sampling location. Other taxa not shown.

### Fungal diversity decreased with increasing urbanization

A noticeable decline in fungal alpha diversity was observed as the sampling locations progressed downstream from non-urban to urban zones. Specifically, a sharp drop-off in diversity was recorded once the urban boundary was crossed. The mean ASV richness ranged from a low of 8 at the furthest downstream location, to a high of 70 at location 3 near the headwater.

ASV richness and Shannon diversity correlated with various urbanization metrics. Low development (e.g., gravel roads, vacant commercial land) was significantly associated with higher fungal richness (P = 0.047) and Shannon diversity (P = 0.099), while cultivated crop cover was significantly associated with lower diversity (P = 0.021 for both diversity measures). Impervious surface area and total developed land surface reduced both richness (P = 0.041) and Shannon diversity (P = 0.011) respectively (Figure 3). Overall, models of land use explained 66% of the variance in fungal richness and 49% of the variance in Shannon diversity (SI Table S2).

**Figure 3.**
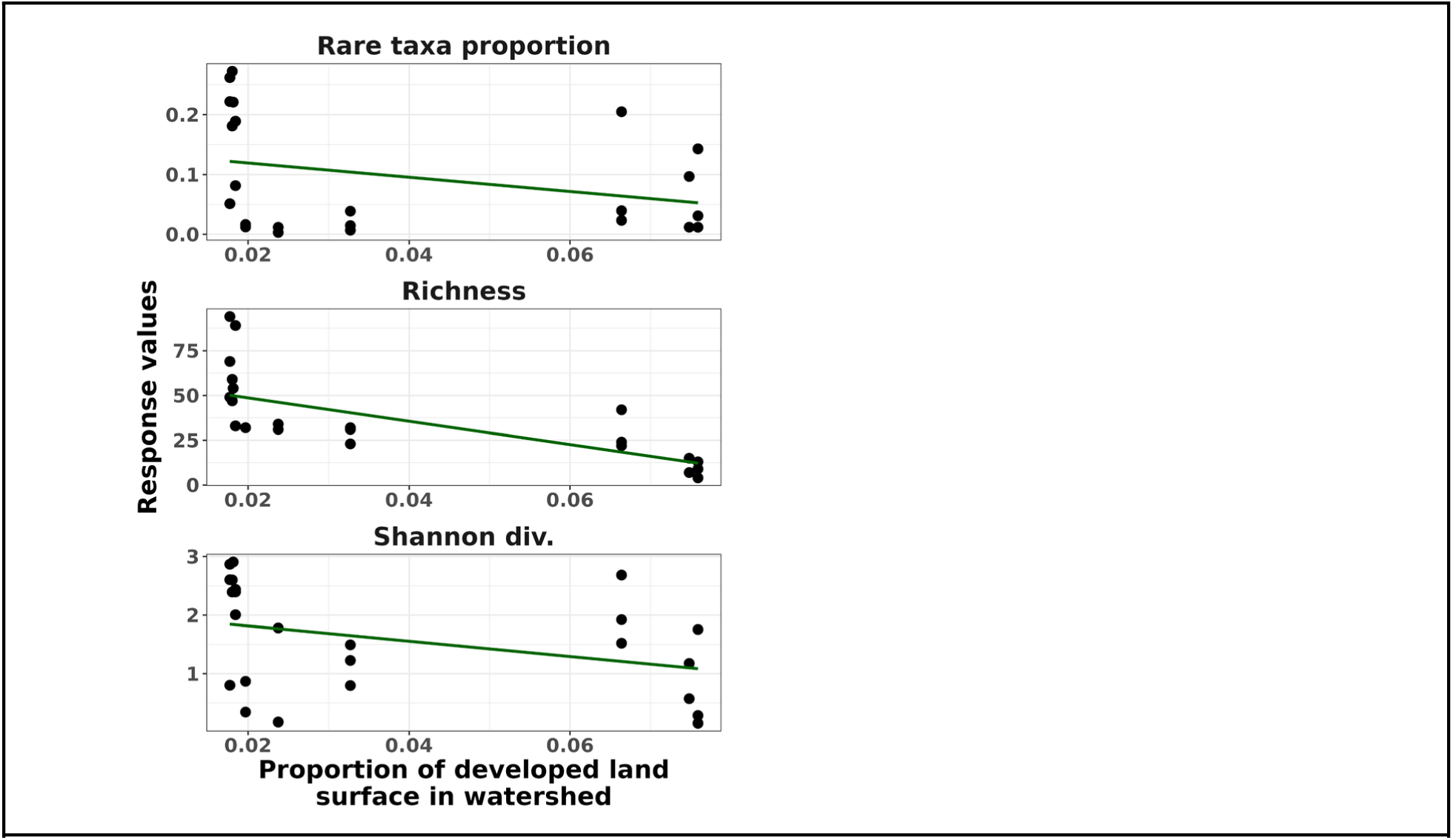
– Regression plots showing the relationship of the proportion of rare taxa (top panel), ASV richness (middle panel), and Shannon diversity (bottom panel) with urbanization. Regression lines were fit with a generalized linear model.

### Rare taxa drove alpha diversity patterns

Rare taxa, defined here as ASVs having less than 0.1% relative abundance in total across all samples, decreased steadily from the headwaters to the urbanized portion of the watershed (P=0.0001) and correlated significantly with the amount of impervious surface area (P=0.003) (SI Table S2). To investigate whether alpha diversity patterns were driven by these rare taxa, we conducted correlation tests and found that the proportion of rare taxa (N rare taxa / N total taxa) at a given site correlated significantly and strongly with both ASV richness (cor = 0.493; P value = 0.014) and Shannon diversity (cor = 0.848; P value < 0.0005) (SI Table S3).

The relative abundance values of the majority of taxa also followed a binomial distribution across samples (SI Figure S2), further suggesting that though communities were dominated by ten highly abundant core fungal ASVs, the remaining community compositions were largely composed of many rare taxa, rather than by semi-abundant taxa. Counts of unique rare taxa in each sample decreased from a high of 78 in the undeveloped headwaters to a low of 1 at the furthest downstream sampling location, with a sharp decline around the transition zone from an undeveloped to an urban-impacted watershed (SI Figure S3).

### Community structure changed with urbanization

Geographic proximity was not a significant predictor of community structure (MRM; P value = 0.55; SI Table S4) even though urbanization metrics covaried with watershed position. However, fungal communities varied significantly along gradients in watershed use. The proportion of developed land (P = 0.006) and proportion of cultivated crops (P = 0.017) in the watershed of a given location together explained 14% of the variance in community structure (S.I. Table S5). A separate PermANOVA model showed that fungal communities were significantly different between the upper non-urbanized and lower urbanized zones of the watershed (P = 0.043; Figure 4).

**Figure 4.**
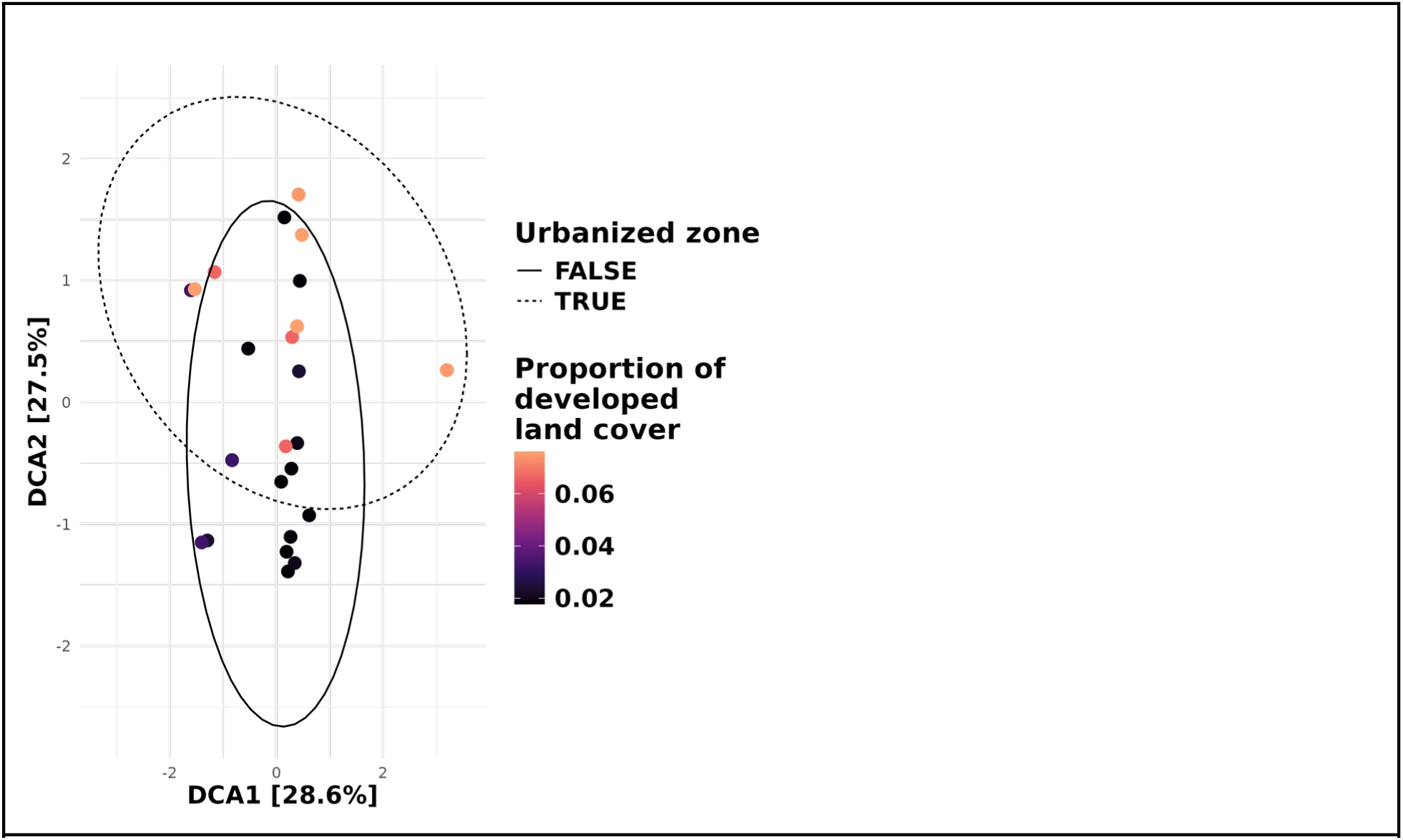
– Detrended correspondence analysis ordination showing Bray-Curtis community dissimilarity among samples. Point color indicates the proportion of developed land area in the cumulative watershed for a given sample. Ellipses represent the 95% confidence intervals around the centroid for samples from locations with highly developed land use (locations 6–10, dotted line) and relatively undeveloped land use (locations 1–5, solid line).

### Fungal guild composition seems unaffected by urbanization

Urbanization and watershed development also had a seemingly significant effect on the balance between mutualist and pathogenic fungi in our model, but there was no consistent pattern between metrics of urbanization. The proportion of ASVs assigned to pathogenic fungal guilds increased with increasing land cover categorized as high development (P = 0.0007), but decreased with overall development. Additionally, the proportion of ASVs assigned to mutualist guilds decreased with increasing overall development (P = 0.024) but increased with the amount of impervious surface. A significant proportion of fungal ASVs could not be reliably assigned to a guild, and this may have affected these models. We are therefore not confident in asserting that urbanization had any affect on the balance of mutualist and pathogen guilds (SI Table S2).

## Discussion

*R. aquatilis* individuals were found consistently throughout the river from our highest point down to to the lowest site. Beyond that site no *R. aquatilis* individuals could be located. Thus, our study represents only a portion of the urbanization gradient along this watershed. Still, we discovered significant changes in fungal diversity and community structure along this gradient. Further, there was a noticeable sudden corresponding with the rapid change in urbanized land cover in the watershed after location 5.

Common heavy metals often associated as urban pollutants (Murray et al. 2004; Li et al. 2012) could not be detected at measurable levels at any point along the river. Metals such as chromium, nickel, copper, zinc, and cadmium have been known to affect many fungi (Abel and Bärlocher 1984; Jones and Muehlchen 1994; Lankinen et al. 2011; Hartikainen et al. 2012; Jin et al. 2018) and so we designed our study to specifically test for those. However, the inability to detect them, and lack of spatial autocorrelation in our results, suggests that other factors associated with urbanization are playing a role in shaping the fungal endophyte communities of aquatic plants. While pH is known to affect aquatic fungi (Rosset and Bärlocher 1985; Dubey et al. 1994), our water samples had highly consistent pH values. Organic pollutants common in urban areas such as pentachlorophenol (Martins et al. 2018) and polycyclic aromatic hydrocarbons (Krauss et al. 2005) are known to affect fungal diversity, but these were not tested for, so it is possible that they may have played a role in shaping the observed patterns.

It is also possible that plant responses to urban stressors could be mediating the fungal communities we observed. We were unable to find any *R. aquatilis* specimens downstream from location 10, and it is possible that these plants were subjected to increasing stress along the urbanization gradient, to the point where no more were able to successfully survive. Plant stress responses can alter endophytic fungal communities via immune signaling and recognition mechanisms (Lu et al. 2021), or by stress attenuating plants’ selective ability on endophytic community assembly (Gao et al. 2020). Transcriptomic analysis of *R. aquatilis* looking for correlations in up-or down-regulation of stress response genes with fungal community could address this hypothesis, but it was not feasible in the current study.

The majority of the fungi in the core mycobiome were known plant pathogens (S.I. Table 1). Our collected plant specimens appeared healthy and had no visible pathogen damage. Some reports list some of these taxa as putatively commensal endophytes however, and it is possible that they were not present in a pathogenic state in our samples. One interesting result is in the widespread presence of *Pseudohalonectria adversaria*, which is generally considered a freshwater saprophytic taxon, but which has been reported to produce nematicidal compounds (Dong et al. 2006). Aquatic *Ranunculus* species have been reported to host pathogenic nematodes (Kohl 2011) and though our methods categorized *P. adversaria* as a saprotroph, it is interesting to consider what role this fungus play in preventing nematode damage in this system. Nematodes remain an understudied component of aquatic plant pathogens (Hong and Moorman 2005) and the interplay between antagonistic fungi and pathogenic nematodes in aquatic plants deserves attention.

Though not a large component of this watershed at any point, cultivated land area did increase along with urbanization metrics such as impermeable surface area. Increased nutrient availability from farm runoff could also alter plants’ dependence on fungal endophytes and nutrient addition has been previously shown to reduce fungal richness in some systems (Henning et al. 2021). To our knowledge, this effect has not been explicitly tested in aquatic plant systems, but it is possible that this could at least partially explain the reduction in fungal endophytic diversity in the lower sites of this watershed.

Taken together, our results show that although aquatic endophyte communities were significantly altered along an urbanization gradient, there is much work ahead to uncover the mechanisms responsible for this observation. Future efforts could productively focus on testing hypotheses regarding organic pollutants, cryptic plant stress condition, and nutrient loads to help understand how aquatic plants in urbanized streams interact with fungi. By the year 2050, urbanization in the US is projected to increase by 88% (Chen et al. 2022) and this is already having strong negative impacts on the health of waterways (Metre et al. 2019). It is crucial that we understand how urbanization alters aquatic plant and fungal interactions in order to more effectively mitigate projected impacts.

### Data availability

Raw sequences generated in this study are available on the Sequence Read Archive under accession **<u>PRJNA1044419</u>**. All metadata and code are available on GitHub at https://github.com/gzahn/Urbanization_and_Endophytes.

## Supplementary Material

**S.I. Figure 1.**
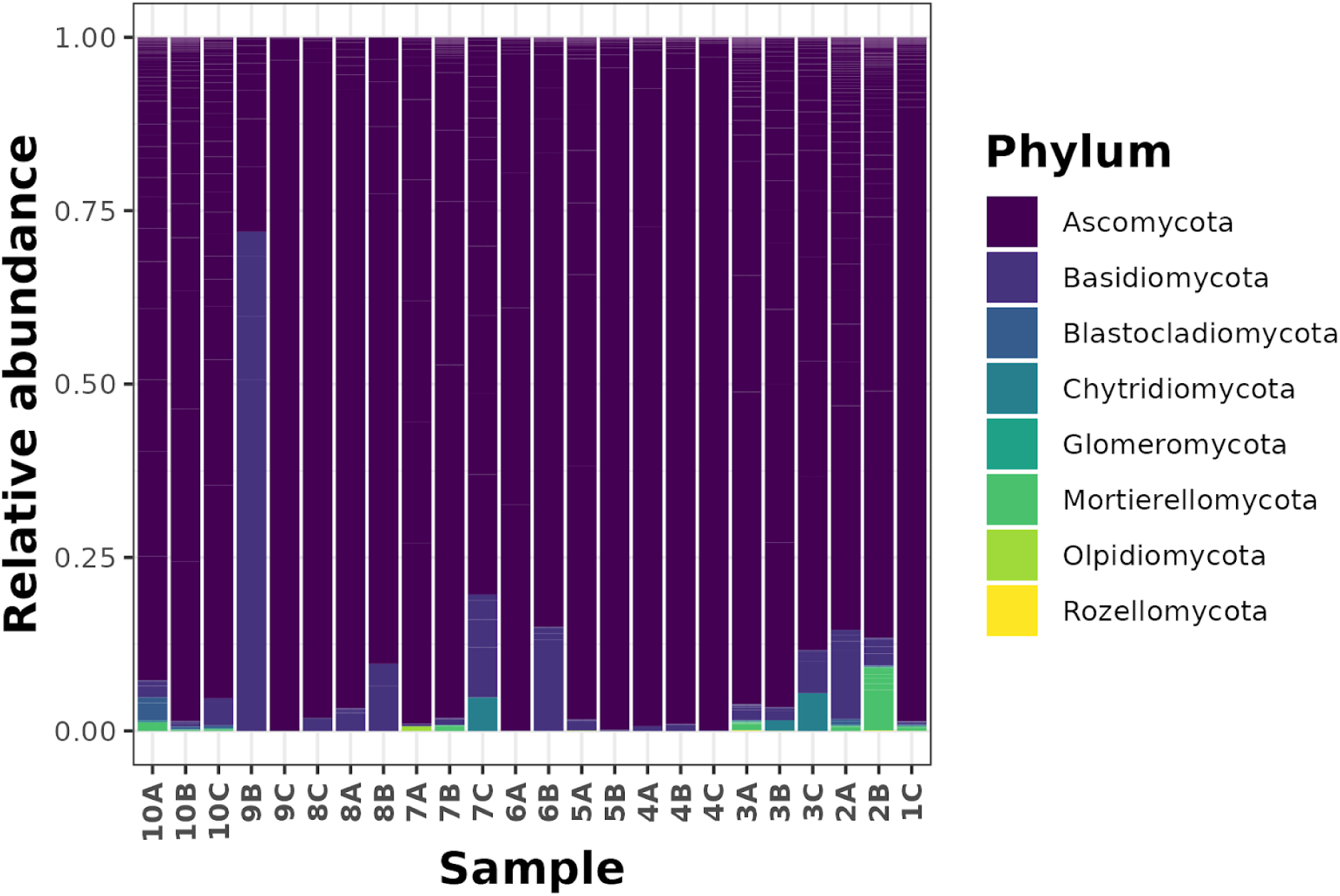
– Stacked bar chart of the relative abundance of ASVs at each sampling location, colored by phylum.

**S.I. Figure 2.**
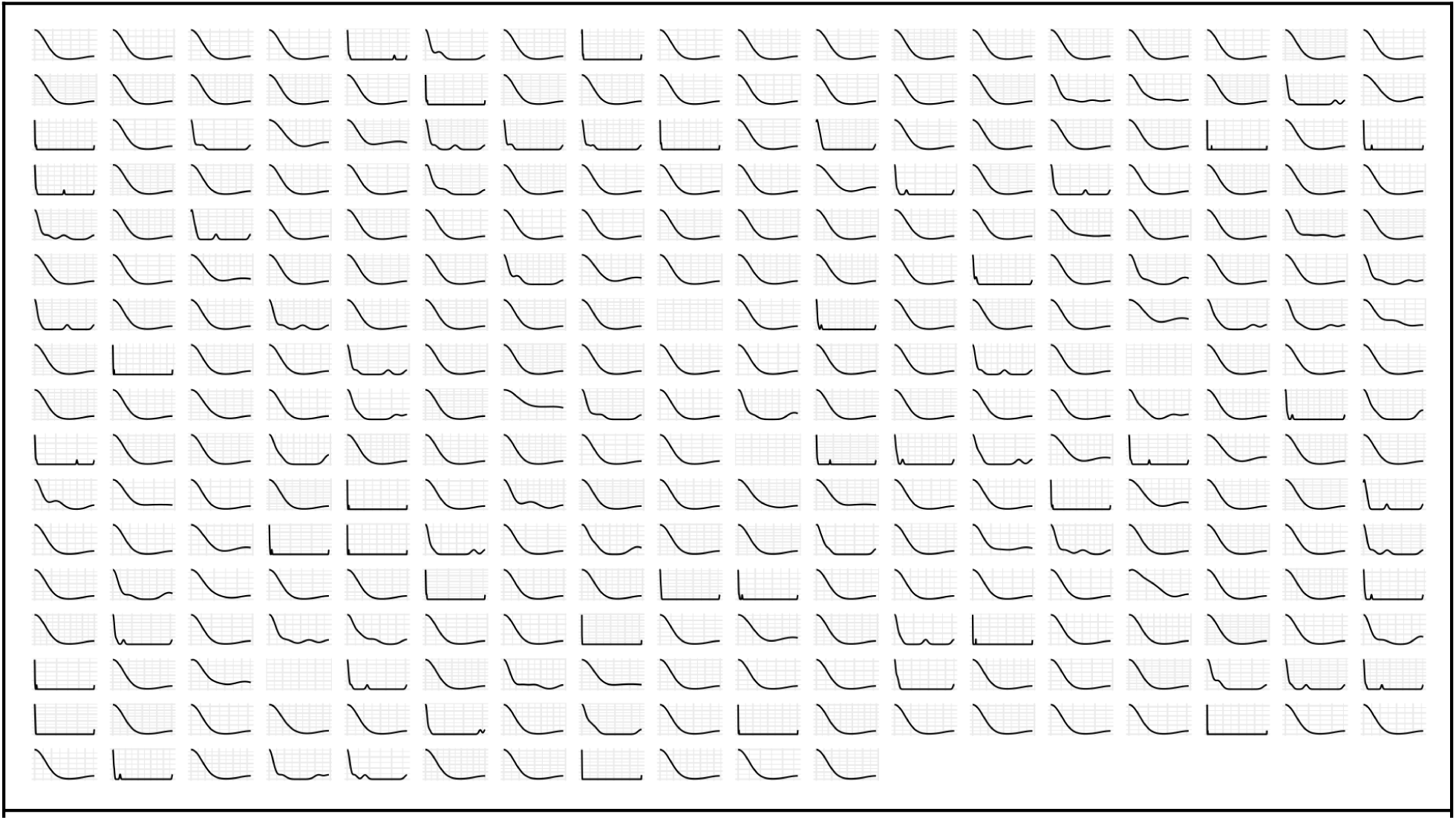
– Relative abundance distributions of all fungal ASVs shown as density plots. X-axis is from 0 to 1. Taxa with no apparent distribution were observed fewer than three times.

**S.I. Figure 3.**
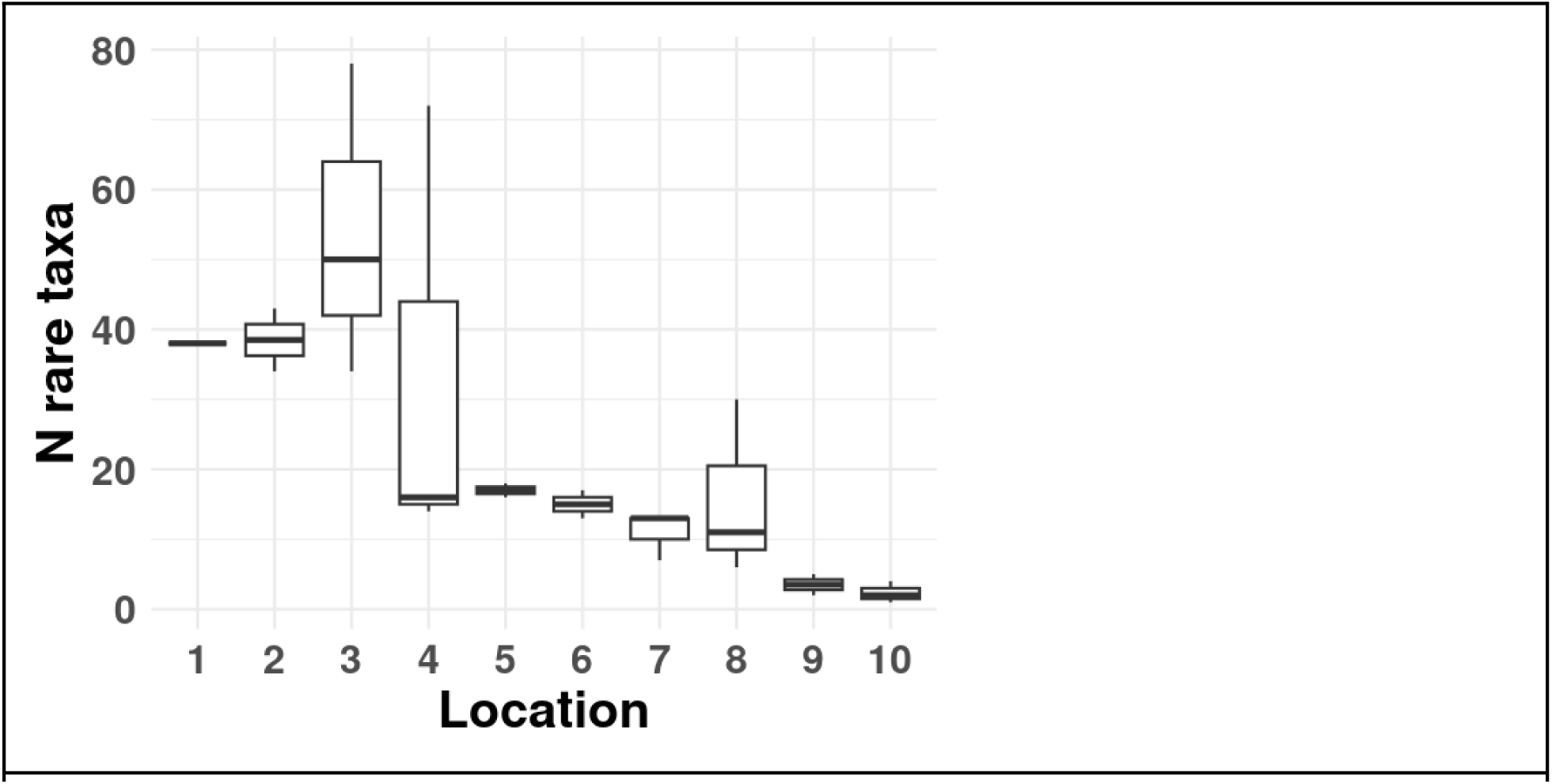
– Numbers of rare taxa observed at each sampling location.

**S.I. Figure 4.**
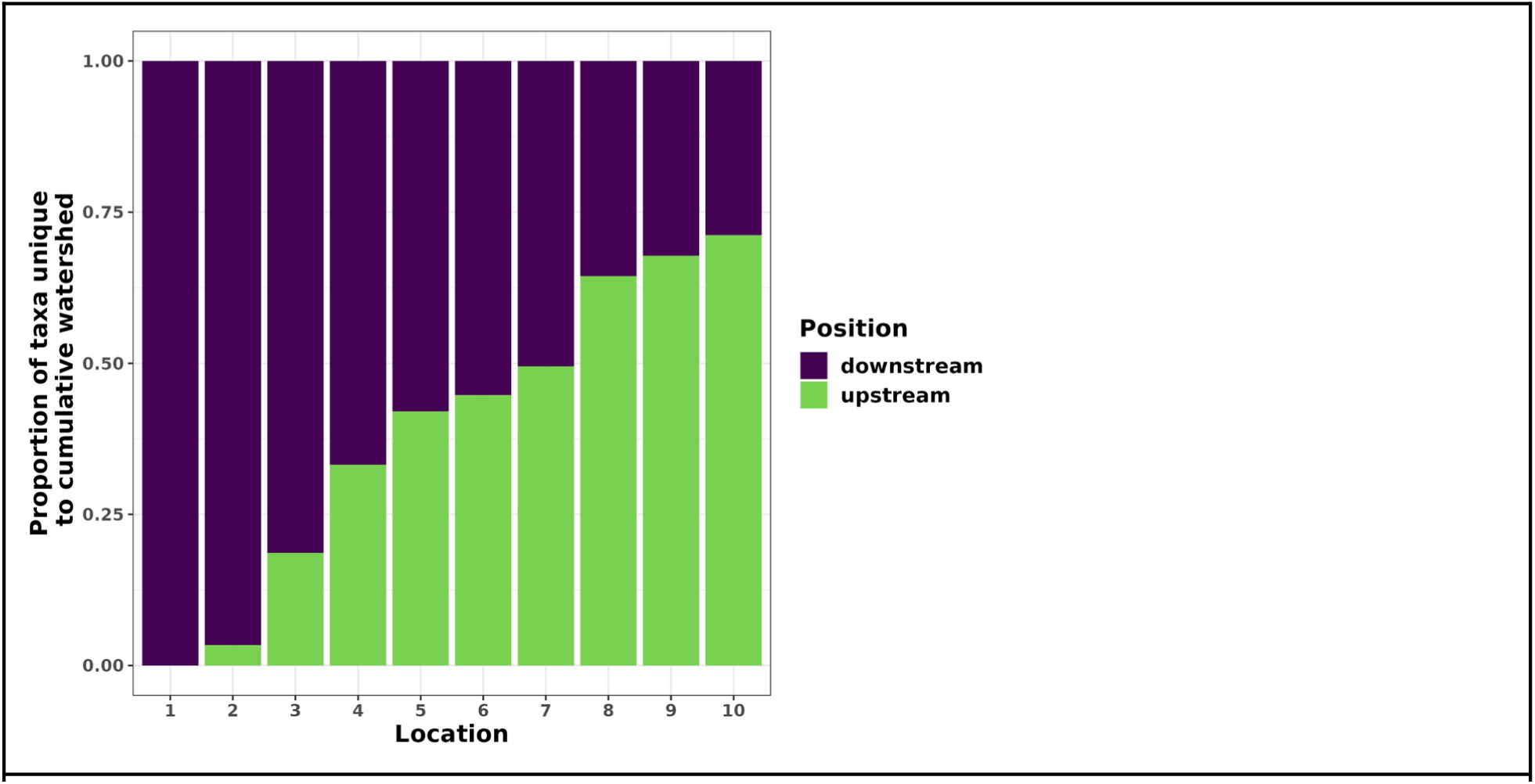
– Proportion of taxa unique to the cumulative upstream (green) and downstream (purple) watershed at a given location, inclusive of that location. X-axis indicates sampling location from 1 (furthest upstream) to 10 (furthest downstream).

**S.I. Figure 5.**
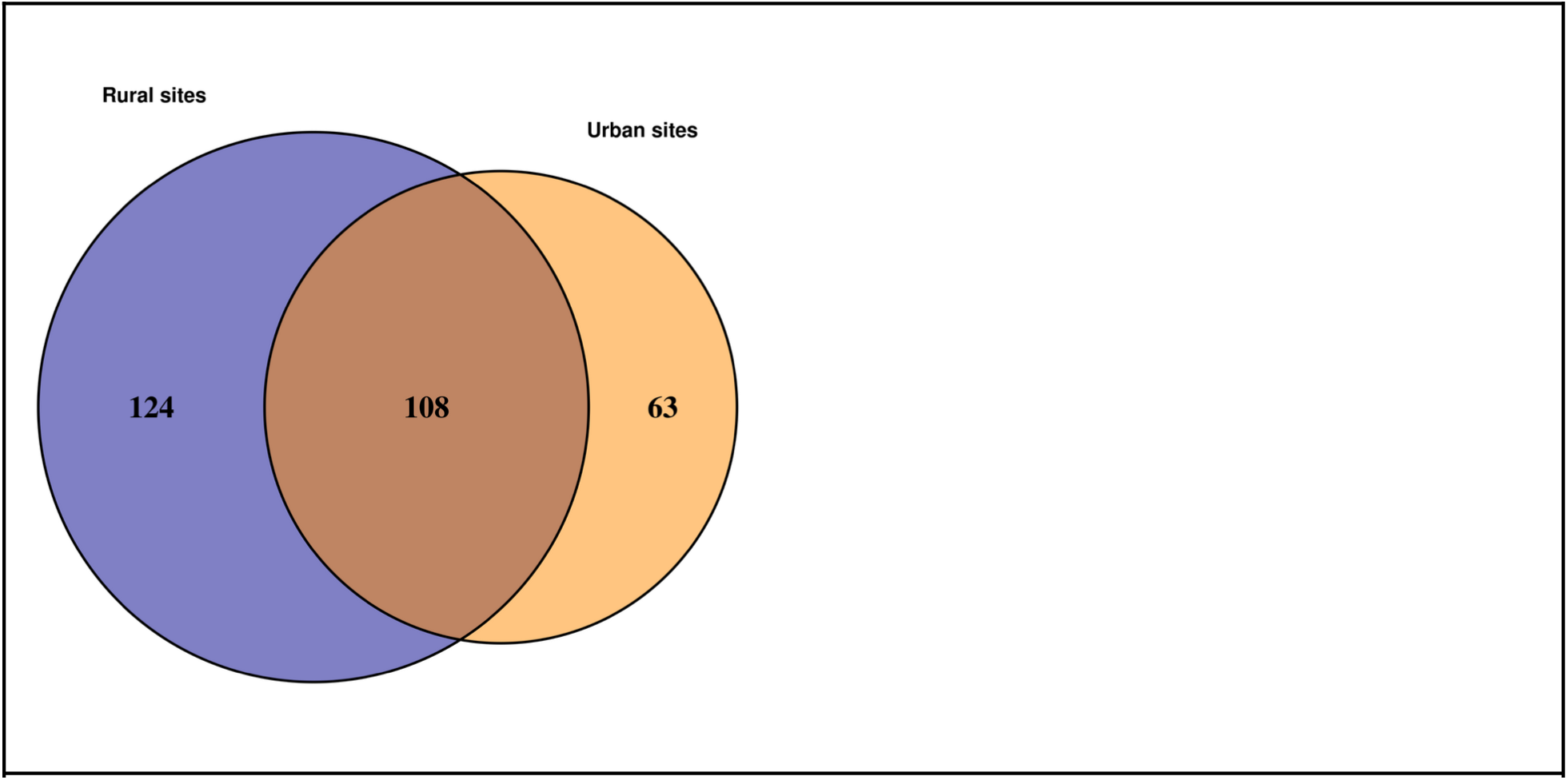
– Venn diagram showing the number of ASVs unique to rural (locations 1-5) and urban sites (impacted by development, locations 6-10), and the number of taxa shared between them.

**SI Table 1.**
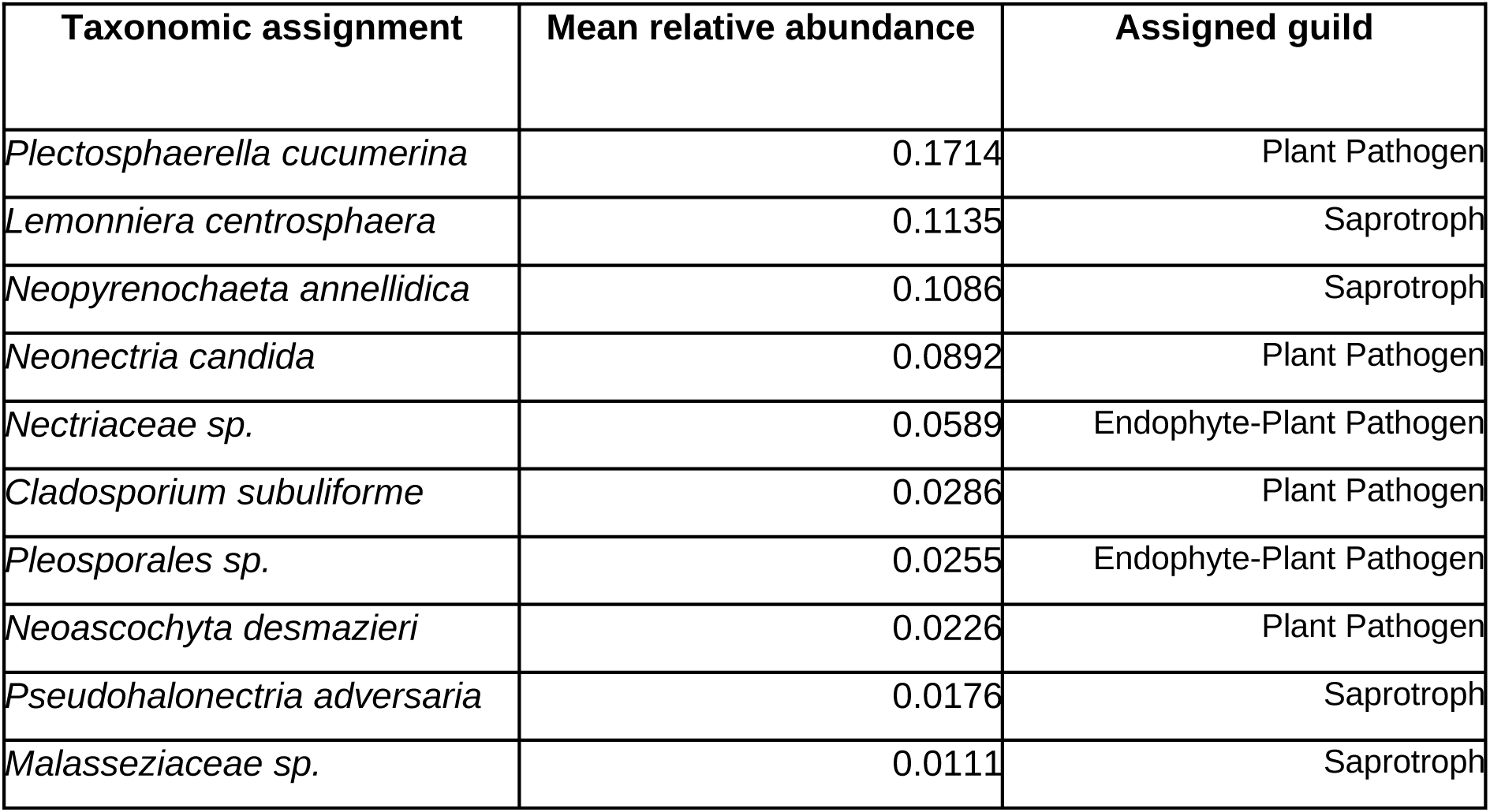
– Assigned taxonomy of the core mycobiome members for the entire study. Core members were defined as ASVs that had greater than 1% relative abundance in at least 50% of all samples. The mean relative abundance and guild assignment of each core mycobiome member is reported.

**S.I. Table 2.**
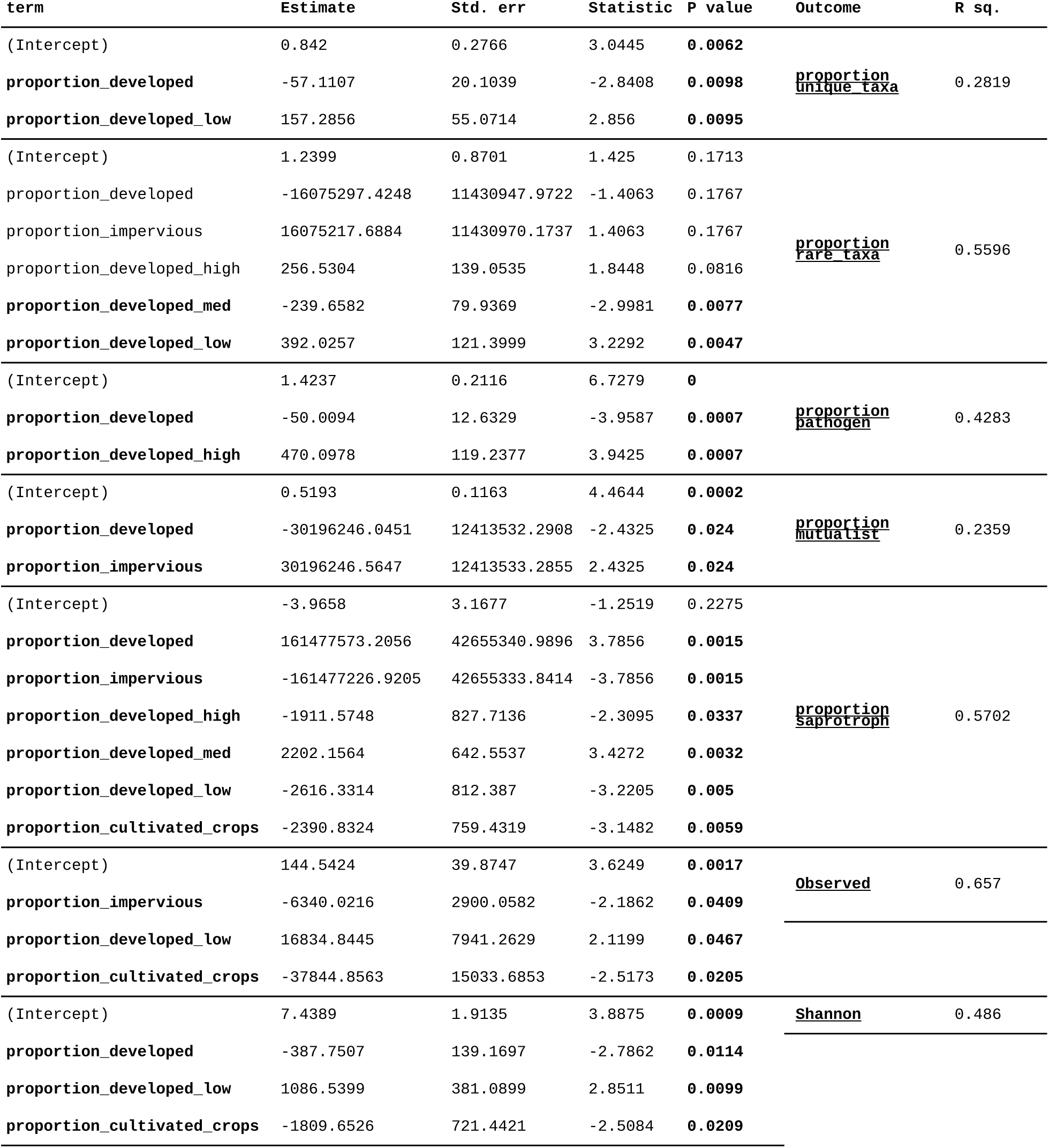
– Diversity and guild composition model results Combined generalized linear regression model results for each tested alpha-diversity outcome (Proportion of rare taxa, richness, and Shannon diversity). Each outcome was modeled independently and refit with the simplified model formulae given by stepwise AIC reduction and including location as an additional predictor. Only the terms from each refit model are shown. P-values < 0.05 are in bold.

**S.I. Table 3.**
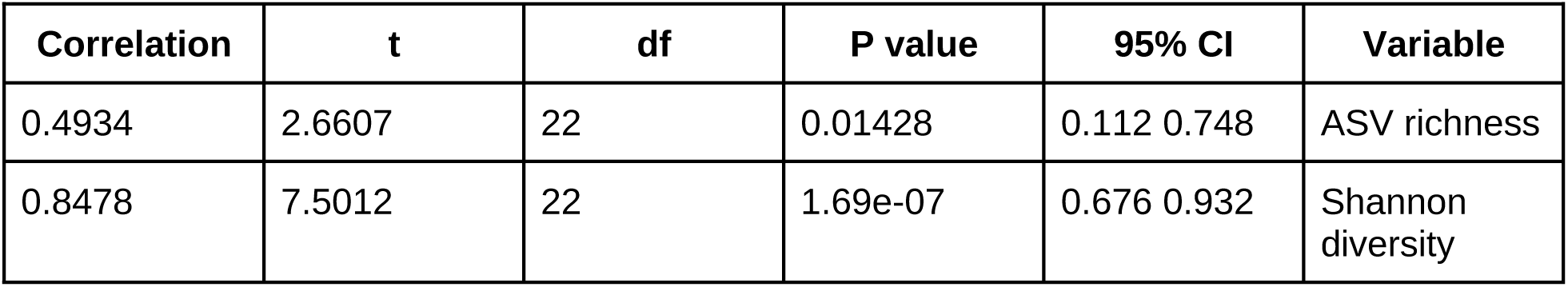
– Correlation test results. Pearson’s product-moment correlation test results. The proportion of rare taxa in a given sample was strongly and significantly correlated with both ASV richness and Shannon diversity.

**S.I. Table 4.**
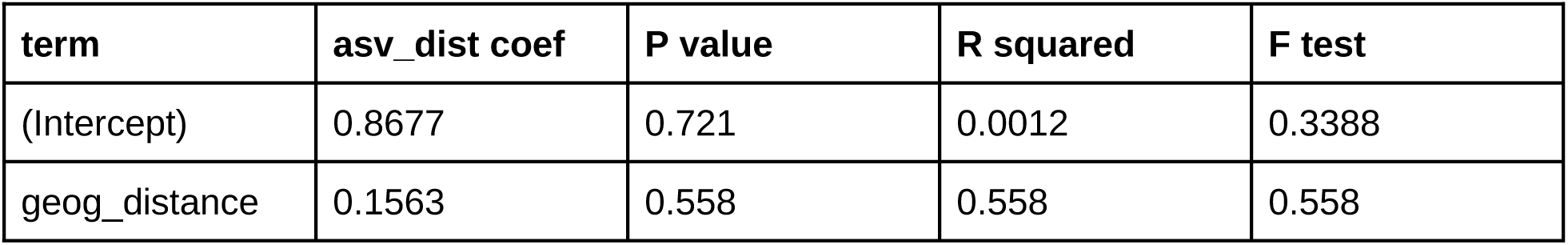
– Multiple Regression on matrices. Regression of Bray-Curtis community dissimilarity matrix on geographic distance matrix. Overall community structure did not significantly correspond to geographic proximity.

**S.I. Table 5.**
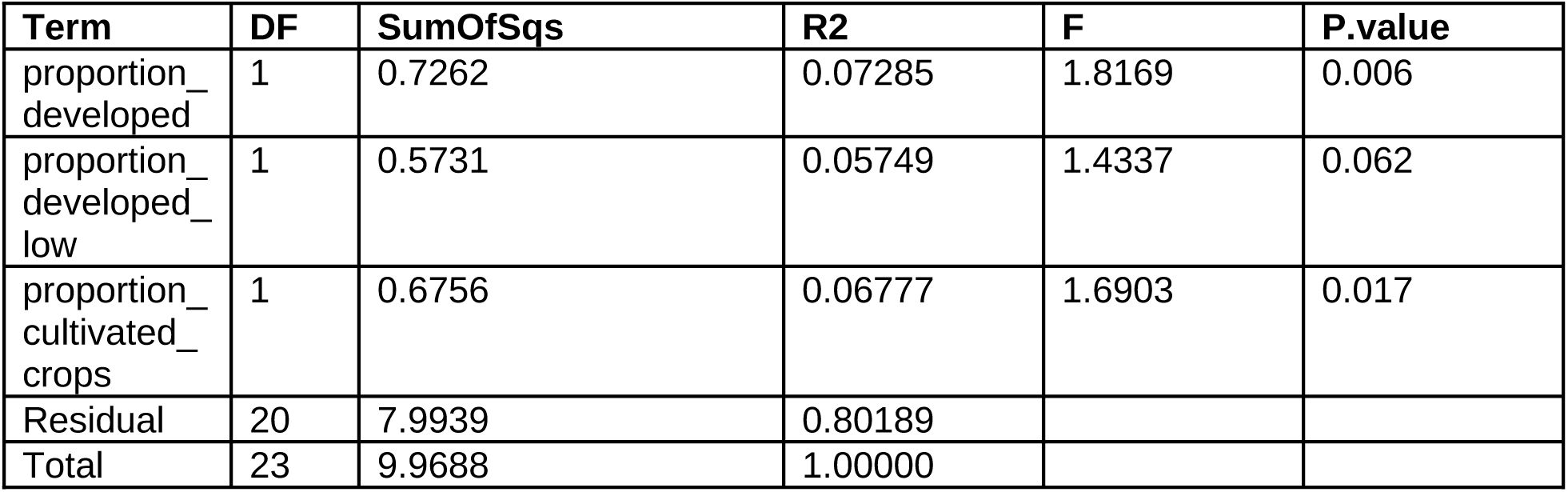
– Permutational ANOVA model results. Terms added sequentially (first to last). Values based on 999 free permutations. P-values < 0.05 are in bold. The proportion of developed or cultivated land in a watershed explained ∼20% of variance in community composition.

**S.I. Table 6.**
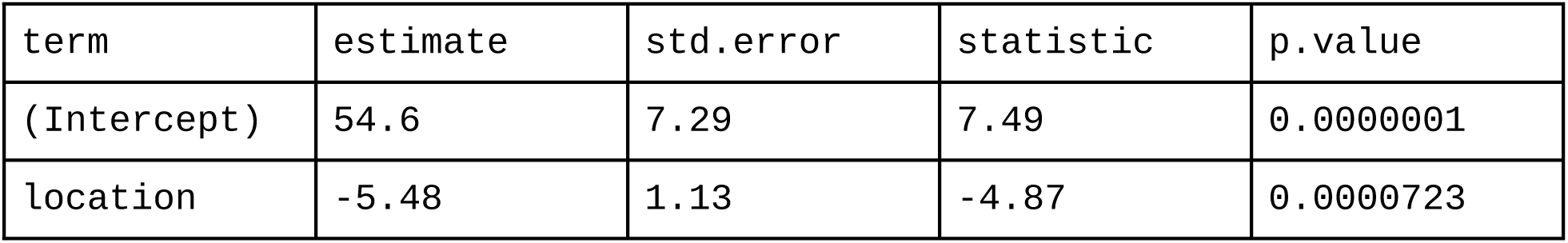
– Counts of rare taxa as a function of downstream location. General linear model.

