## supplemental tables and figures for "Aquatic plant mycobiomes change with watershed urbanization"

**Supplementary Material**

**S.I. Figure 1** - Stacked bar chart of the relative abundance of ASVs at each sampling location, colored by phylum.

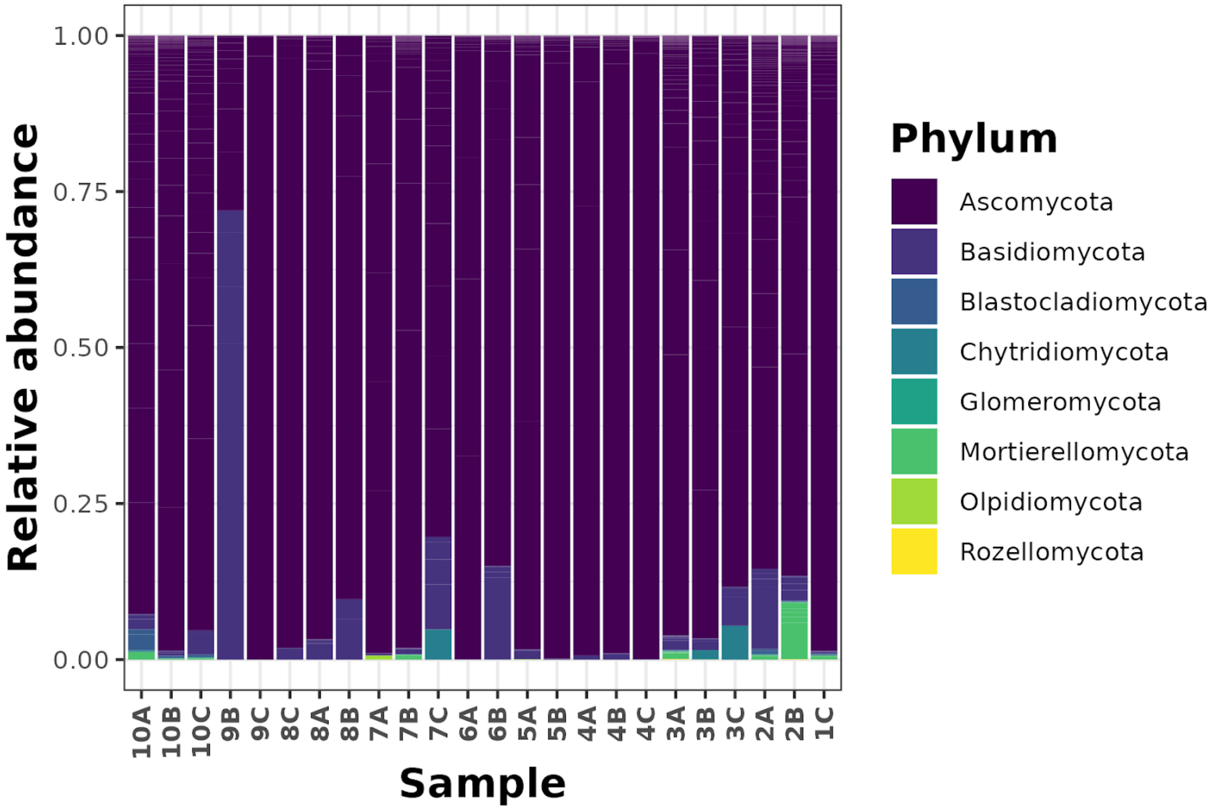

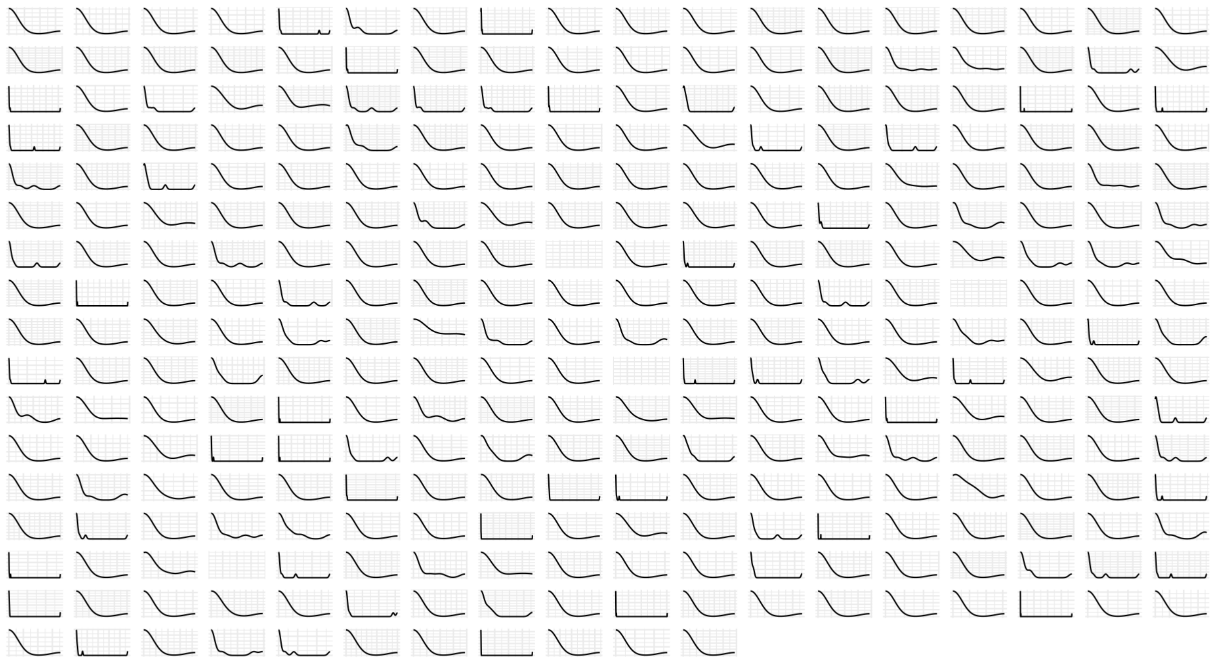

**S.I. Figure 2** - Relative abundance distributions of all fungal ASVs shown as density plots. X-axis is from 0 to 1. Taxa with no apparent distribution were observed fewer than three times.

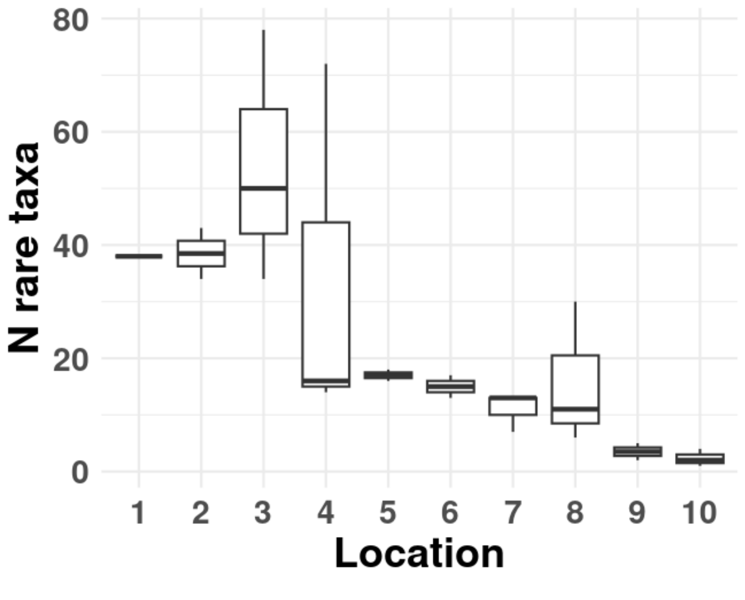


**S.I. Figure 3 -** Numbers of rare taxa observed at each sampling location.

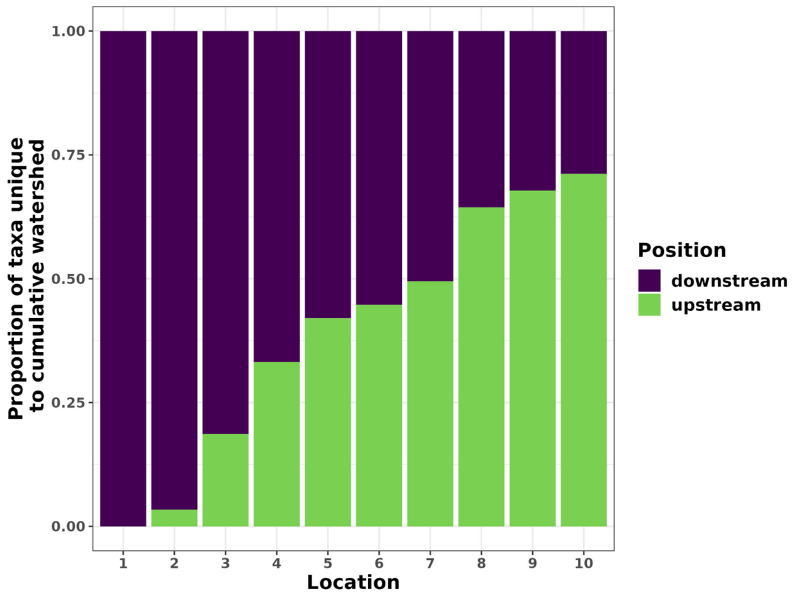


**S.I. Figure 4** - Proportion of taxa unique to the cumulative upstream (green) and downstream (purple) watershed at a given location, inclusive of that location. X-axis indicates sampling location from 1 (furthest upstream) to 10 (furthest downstream).

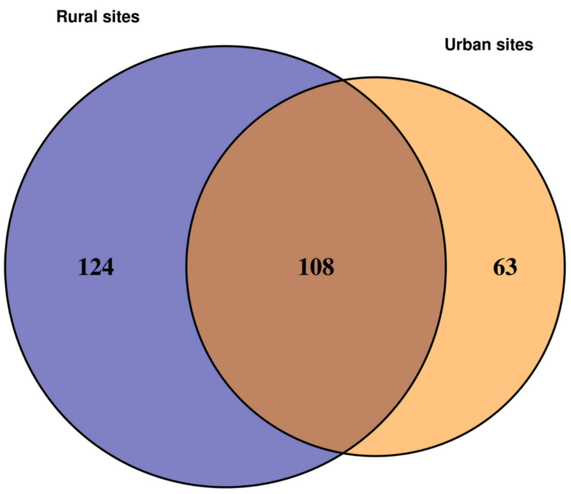


**S.I. Figure 5** - Venn diagram showing the number of ASVs unique to rural (locations 1-5) and urban sites (impacted by development, locations 6-10), and the number of taxa shared between them.

| Taxonomic assignment | Mean relative abundance | Assigned guild |
|---|---|---|

| | | |
|---|---|---|
| *Plectosphaerella cucumerina* | 0.1714 | Plant Pathogen |
| *Lemonniera centrosphaera* | 0.1135 | Saprotroph |
| *Neopyrenochaeta annellidica* | 0.1086 | Saprotroph |
| *Neonectria candida* | 0.0892 | Plant Pathogen |
| *Nectriaceae sp.* | 0.0589 | Endophyte-Plant Pathogen |
| *Cladosporium subuliforme* | 0.0286 | Plant Pathogen |
| *Pleosporales sp.* | 0.0255 | Endophyte-Plant Pathogen |
| *Neoascochyta desmazieri* | 0.0226 | Plant Pathogen |
| *Pseudohalonectria adversaria* | 0.0176 | Saprotroph |
| *Malasseziaceae sp.* | 0.0111 | Saprotroph |

**SI Table 1 -** Assigned taxonomy of the core mycobiome members for the entire study. Core members were defined as ASVs that had greater than 1% relative abundance in at least 50% of all samples. The mean relative abundance and guild assignment of each core mycobiome member is reported.

**S.I. Table 2 - Diversity and guild composition model results**

Combined generalized linear regression model results for each tested alpha-diversity outcome (Proportion of rare taxa, richness, and Shannon diversity). Each outcome was modeled independently and refit with the simplified model formulae given by stepwise AIC reduction and including location as an additional predictor. Only the terms from each refit model are shown. P-values < 0.05 are in bold.

| term | Estimate | Std. err | Statistic | P value | Outcome | R sq. |
|---|---|---|---|---|---|---|
| (Intercept) | 0.842 | 0.2766 | 3.0445 | **0.0062** | | |
| **proportion_developed** | -57.1107 | 20.1039 | -2.8408 | **0.0098** | proportion unique_taxa | 0.2819 |
| **proportion_developed_low** | 157.2856 | 55.0714 | 2.856 | **0.0095** | | |
| (Intercept) | 1.2399 | 0.8701 | 1.425 | 0.1713 | | |
| proportion_developed | -16075297.4248 | 11430947.9722 | -1.4063 | 0.1767 | | |
| proportion_impervious | 16075217.6884 | 11430970.1737 | 1.4063 | 0.1767 | | |
| proportion_developed_high | 256.5304 | 139.0535 | 1.8448 | 0.0816 | proportion rare_taxa | 0.5596 |
| **proportion_developed_med** | -239.6582 | 79.9369 | -2.9981 | **0.0077** | | |
| **proportion_developed_low** | 392.0257 | 121.3999 | 3.2292 | **0.0047** | | |
| (Intercept) | 1.4237 | 0.2116 | 6.7279 | **0** | | |
| **proportion_developed** | -50.0094 | 12.6329 | -3.9587 | **0.0007** | proportion pathogen | 0.4283 |
| **proportion_developed_high** | 470.0978 | 119.2377 | 3.9425 | **0.0007** | | |
| (Intercept) | 0.5193 | 0.1163 | 4.4644 | **0.0002** | | |
| **proportion_developed** | -30196246.0451 | 12413532.2908 | -2.4325 | **0.024** | proportion mutualist | 0.2359 |
| **proportion_impervious** | 30196246.5647 | 12413533.2855 | 2.4325 | **0.024** | | |
| (Intercept) | -3.9658 | 3.1677 | -1.2519 | 0.2275 | | |
| **proportion_developed** | 161477573.2056 | 42655340.9896 | 3.7856 | **0.0015** | | |
| **proportion_impervious** | -161477226.9205 | 42655333.8414 | -3.7856 | **0.0015** | | |
| **proportion_developed_high** | -1911.5748 | 827.7136 | -2.3095 | **0.0337** | proportion saprotroph | 0.5702 |
| **proportion_developed_med** | 2202.1564 | 642.5537 | 3.4272 | **0.0032** | | |
| **proportion_developed_low** | -2616.3314 | 812.387 | -3.2205 | **0.005** | | |
| **proportion_cultivated_crops** | -2390.8324 | 759.4319 | -3.1482 | **0.0059** | | |
| (Intercept) | 144.5424 | 39.8747 | 3.6249 | **0.0017** | | |
| **proportion_impervious** | -6340.0216 | 2900.0582 | -2.1862 | **0.0409** | Observed | 0.657 |
| **proportion_developed_low** | 16834.8445 | 7941.2629 | 2.1199 | **0.0467** | | |
| **proportion_cultivated_crops** | -37844.8563 | 15033.6853 | -2.5173 | **0.0205** | | |
| (Intercept) | 7.4389 | 1.9135 | 3.8875 | **0.0009** | | |
| **proportion_developed** | -387.7507 | 139.1697 | -2.7862 | **0.0114** | Shannon | 0.486 |
| **proportion_developed_low** | 1086.5399 | 381.0899 | 2.8511 | **0.0099** | | |
| **proportion_cultivated_crops** | -1809.6526 | 721.4421 | -2.5084 | **0.0209** | | |

**S.I. Table 3 - Correlation test results.**
Pearson's product-moment correlation test results. The proportion of rare taxa in a given sample was strongly and significantly correlated with both ASV richness and Shannon diversity.

| Correlation | t | df | P value | 95% CI | Variable |
|---|---|---|---|---|---|
| 0.4934 | 2.6607 | 22 | 0.01428 | 0.112 0.748 | ASV richness |
| 0.8478 | 7.5012 | 22 | 1.69e-07 | 0.676 0.932 | Shannon diversity |

**S.I. Table 4 - Multiple Regression on matrices.** Regression of Bray-Curtis community dissimilarity matrix on geographic distance matrix. Overall community structure did not significantly correspond to geographic proximity.

| **term** | **asv_dist coef** | **P value** | **R squared** | **F test** |
|---|---|---|---|---|
| (Intercept) | 0.8677 | 0.721 | 0.0012 | 0.3388 |
| geog_distance | 0.1563 | 0.558 | 0.558 | 0.558 |

**S.I. Table 5 -** Permutational ANOVA model results. Terms added sequentially (first to last). Values based on 999 free permutations. P-values < 0.05 are in bold. The proportion of developed or cultivated land in a watershed explained ~20% of variance in community composition.

| **Term** | **DF** | **SumOfSqs** | **R2** | **F** | **P.value** |
|---|---|---|---|---|---|
| proportion_developed | 1 | 0.7262 | 0.07285 | 1.8169 | 0.006 |
| proportion_developed_low | 1 | 0.5731 | 0.05749 | 1.4337 | 0.062 |
| proportion_cultivated_crops | 1 | 0.6756 | 0.06777 | 1.6903 | 0.017 |
| Residual | 20 | 7.9939 | 0.80189 | | |
| Total | 23 | 9.9688 | 1.00000 | | |

**S.I. Table 6 -** Counts of rare taxa as a function of downstream location. General linear model.

| term | estimate | std.error | statistic | p.value |
|---|---|---|---|---|
| (Intercept) | 54.6 | 7.29 | 7.49 | 0.0000001 |
| location | -5.48 | 1.13 | -4.87 | 0.0000723 |
